# An Assay for Apoptosis detection based on Quantification of Multi nuclei feature and Nucleus to Cytoplasm ratio in *S. cerevisiae* cells treated with Acetic Acid and Hydrogen peroxide

**DOI:** 10.1101/2020.03.11.987024

**Authors:** Narendra K Bairwa, Heena Shoket, Monika Pandita, Meenu Sharma

## Abstract

The programmed cell death, apoptosis is a complex universal biological process in all types of eukaryotes ranging from single cell to multi-cellular organisms. The markers for apoptosis have been studied by assays based on both biochemical as well as microscopy however most assays are not affordable for many smaller labs. Acetic acid and hydrogen peroxide both induce apoptosis at higher concentrations in *S. cerevisiae.* Here we describe an assay system for the detection of apoptosis features based on DAPI staining followed by fluorescence microscopy in the cells treated with apoptosis inducing concentration of acetic acid and hydrogen peroxide. In this assay both untreated and cells treated with acetic acid and hydrogen peroxide were stained with DAPI and observed for the late stage apoptosis feature, Nuclear DNA fragmentation based multi nuclei centers and increase in the nuclear region enlargement. Further the multi nuclei feature and enlarged nuclei region as nucleus to cytoplasm ratio was quantified using Image J software. We report that *S. cerevisiae* strain BY4741 cells when treated with apoptosis inducing doses of acetic acid (140mM) and hydrogen peroxide (10mM) for 200 minutes, showed apoptosis marker feature such as nuclear region enlargement with multi-nuclei feature due to nuclear DNA fragmentation and increased nucleus to cytoplasm ratio when compared with untreated cells. We propose that this assay can be utilized for scoring the quantitative apoptotic feature as increase in multi-nuclei centers due to DNA fragmentation and nucleus to cytoplasm ratio as an indicator of apoptosis in *S. cerevisiae* upon treatment with apoptosis inducing agents. The assay system described here is easy to perform and affordable for the smaller lab to analyze the apoptotic features in *S. cerevisiae* cells which can be applied to other system as well.

## Introduction

Cellular suicide, termed as apoptosis is critical process of removing damaged or unwanted cells in multi-cellular organisms and was first described in animals (Kerr *et al.* 1972; Green 2005). The apoptosis process is important during normal ageing and in the containment of the infection caused by bacteria and viral agent. The deregulation of the apoptosis process in multi-cellular organism, especially human may result in diseases such as cancer, neuro-degeneration and viral infections. Cells undergoing apoptosis displays various marker features such as phosphatidylserine externalization on outer membrane, membrane blebbing, protein leakage from the mitochondria, increased caspase activity, chromatin condensation and DNA fragmentation (Green 2005; Elmore 2007). The Nuclear DNA fragmentation is a key feature of the cells which undergo apoptosis in the later stages of cell death. *Saccharomyces cerevisiae* has been used as a model organism for studying apoptosis as it shares the conserved pathways with mammalian cells (Madeo *et al.* 2002). *S. cerevisia*e has particularly been used in elucidation of apoptosis pathways induced by acetic acid. Acetic acid is a by-product of alcoholic fermentation in yeast and affect the bio-ethanol production in industrial fermentation (Liu and Blaschek 2010). Acetic acid also acts as an apoptotic agent at higher concentrations (Lee *et al.* 2011) and has cytotoxic effect which has been utilized by food industry as preservative at higher concentration. However at lower concentration of 0.2 to 0.6 g/l, it serves as a carbon and energy source for *S. cerevisiae* (Sousa *et al.* 2012a; Sousa *et al.* 2012b).The growing *S. cerevisiae* cells undergo apoptosis when treated with 80mM acetic acid for 200 minutes (Ludovico *et al.* 2001; Giannattasio *et al.* 2005) and exposure to lethal concentrations of 140mM acetic acid lead to cell death (Rego *et al.* 2014b). Hydrogen peroxide is one of the first oxidative stress agent known to induce apoptosis in yeast. The hydrogen peroxide at the concentration of 10mM is also used as an apoptotic agent for several cell types including cell lines and tumour (Xiang *et al.* 2016). Several assays which have been used for studying the mammalian cell apoptosis, extended to yeast cells for studying the apoptosis, including, nuclear DNA fragmentation (TUNEL Assay), exposure of phosphatidylserine to the cytoplasmic leaflet (Annexin V Staining) and release of cytochrome C from mitochondria (Hardwick and Cheng 2004). However most of the methods for apoptosis detection are complex and costly except CFU counting and bright field microscopy assay which are indirect. Here we describe a easy to perform method to score the apoptosis features which combined the two major observations first, multi-nuclei phenotype due to nuclear migration defects, earlier described by(Brachat *et al.* 1998) for studying spindle pole body defects second, increased nucleus to cytoplasm ratio due to nuclear DNA fragmentation mediated nuclear region enlargement *in vivo* in *S. cerevisiae cells* when treated with apoptosis inducing concentration of the acetic acid and hydrogen peroxide. Both treated and untreated cells were stained with DAPI and analyzed for both the apoptosis features. The semi-quantitative growth assay was also carried out for comparative growth analysis between treated and untreated cells. We report that cells undergoing apoptosis exhibits both increased in the multi nuclei phenotype and nucleus to cytoplasm ratio which can be performed routinely in smaller lab without the use of expensive biochemical to analyze the apoptosis in *S. cerevisiae* cells.

## Materials& Methods

1. Yeast strain: BY4741 (*MATa his3Δ1 leu2Δ0 met15Δ0 ura3Δ0*)
2. YPD liquid medium: 1% Yeast extract, 2% Peptone, 2% Glucose.
3. Flasks, for yeast culture (Autoclaved)
4. Acetic acid (stock solution - 17.4 Molar)
5. Hydrogen peroxide (Stock solution – 1.76 Molar)
6. DAPI staining solution (Sigma)
7. Leica Fluorescence microscope DM3000 with multi-pass filter sets specific for viewing DAPI stained cells.
8. Micro slides and micro cover slips

### Method

1. WT strain BY4741 was streaked out from -20°C glycerol stock onto YPD agar plates and incubated at 30°C for 2 days in Thermo Scientific incubator.
2. From YPD agar plate, single colony was picked and inoculated into autoclaved fresh 10ml YPD broth and incubated at 30°C overnight at 200 rpm in Thermo Scientific incubator shaker.
3. Next day, overnight grown culture was inoculated into fresh 10ml YPD medium in 1:20 ratio with or without 140mM Acetic acid (OD_600nm_ = 0.2) and 10mM H_2_O_2_ and incubated for 200 minutes until mid-log phase at 200 rpm.
4. Both untreated and treated cells were harvested by centrifugation at 2000 rpm for 2-3 minutes and washed with distilled water. An aliquot of cells from both untreated and treated were adjusted equally by optical density measurement at 600nM and 10 fold serially diluted. The serially diluted cells were spotted on YPD plate and incubated at 30°C for 24-36 hrs for analysis of semi-quantitative growth after apoptosis using doses of acetic acid and hydrogen peroxide treatment.
5. Further an aliquot of cells was suspended in 1X Phosphate Buffer saline (PBS) and fixation was carried out by adding 70% ethanol for 10 minutes and centrifuged for 2.5 minutes at 2500 rpm.
6. DAPI stain (1mg/ml) to final concentration (2.5µg/ml) was added and incubated for 5 minutes at room temperature and visualized under UV light of fluorescence microscope with 100X magnification. Images were acquired for analysis of the nucleus to cytoplasm ratio and multi nuclei phenotype in both the untreated and treated cells.
7. For quantification of the nucleus to cytoplasm ratio, Image J (http://imagej.nih.gov) software was used. The image acquired was analysed using Image J toolbox. The scale tool box was used to measure the area of the nucleus and cytoplasm separately of the DAPI stained cell **(Figure 1 B**). A total of 25 DAPI stained mother cells from each treated and untreated were analysed for calculation of area of nucleus and cytoplasm. The average of the area of nucleus of 25 cells was divided by the average of the area of cytoplasm of 25 cells and a ratio was drawn for both treated and untreated cells and plotted as a bar diagram to determine the difference between treated and untreated.
8. For quantification of increased multi nuclei phenotype due to nuclear DNA fragmentation and migration defects, 100 DAPI stained cells were observed and categorized on the basis of presence of nuclei within the cell as 0, 1, 2 or >3 nuclei per cell. While 1 or 2 nuclei represent the normal cell phenotype (**Figure 2A**), presence of more than 2 nuclei in a cell is due to DNA fragmentation indicated as apoptotic feature or nuclear defects. The percentage of the cells showed 1 nucleus, 2 nuclei, and multi nuclei was counted and plotted as bar diagram.

## Results & Discussion

In this assay, *S. cerevisiae* laboratory strain BY4741 was used for apoptosis feature analysis after treatment with the lethal dose of acetic acid (140 mM) and hydrogen peroxide (10 mM) for 200 minutes. Acetic acid is an apoptosis inducing agent and exposure to lethal concentrations of 140 mM causes cell death (Rego *et al.* 2014a). After treatment, the cells were fixed and stained with 4’, 6-diamidino-2-phenylindole (DAPI) stain and analysed by microscopy. The untreated cells showed the compact nucleus when stained with DAPI (**Figure 1A, E**) however cells treated with acetic acid and hydrogen peroxide showed the nuclear DNA fragmentation with increase in the nuclear chromatin region (**Figure 1A, E**) as measured using Image J software in both treated and untreated cells (**Figure 1B**). The changes in the nuclear area occurred due to fragmentation of the chromatin, which is the hallmark of apoptosis (Green 2005).We further tested the impact of acetic acid and hydrogen peroxide treatment on the cellular growth by spot assays. The acetic acid and hydrogen peroxide treated and untreated cells, both were serially diluted and spotted on YPD+ agar plate and incubated at 30° C for 24 hrs and relative growth was analysed. The treated cells showed the growth defect in comparison to the untreated cells (**Figure 1D, G**) indicating the effectiveness of the treatment. Next, we quantified the nucleus to cytoplasm ratio by measuring the nucleus and the cytoplasm area using the Image J software. We analysed nucleus region and the cytoplasm area including nuclear region of 25 DAPI stained mother cells on the acquired images by Image J software. The average of nucleus area and the cytoplasm area was calculated and ratio of nucleus to cytoplasm was drawn as mentioned in (**Figure 1B**). It was observed that the apoptosis inducing concentration of acetic acid and hydrogen peroxide caused apoptosis phenotype such as nuclear fragmentation (dispersed nuclei) and chromatin condensation (incompletely rounded nuclei) and 2-fold increase in the nucleus to cytoplasm ratio in comparison to untreated cells (**Figure 1C, F**).

**Figure 1.**
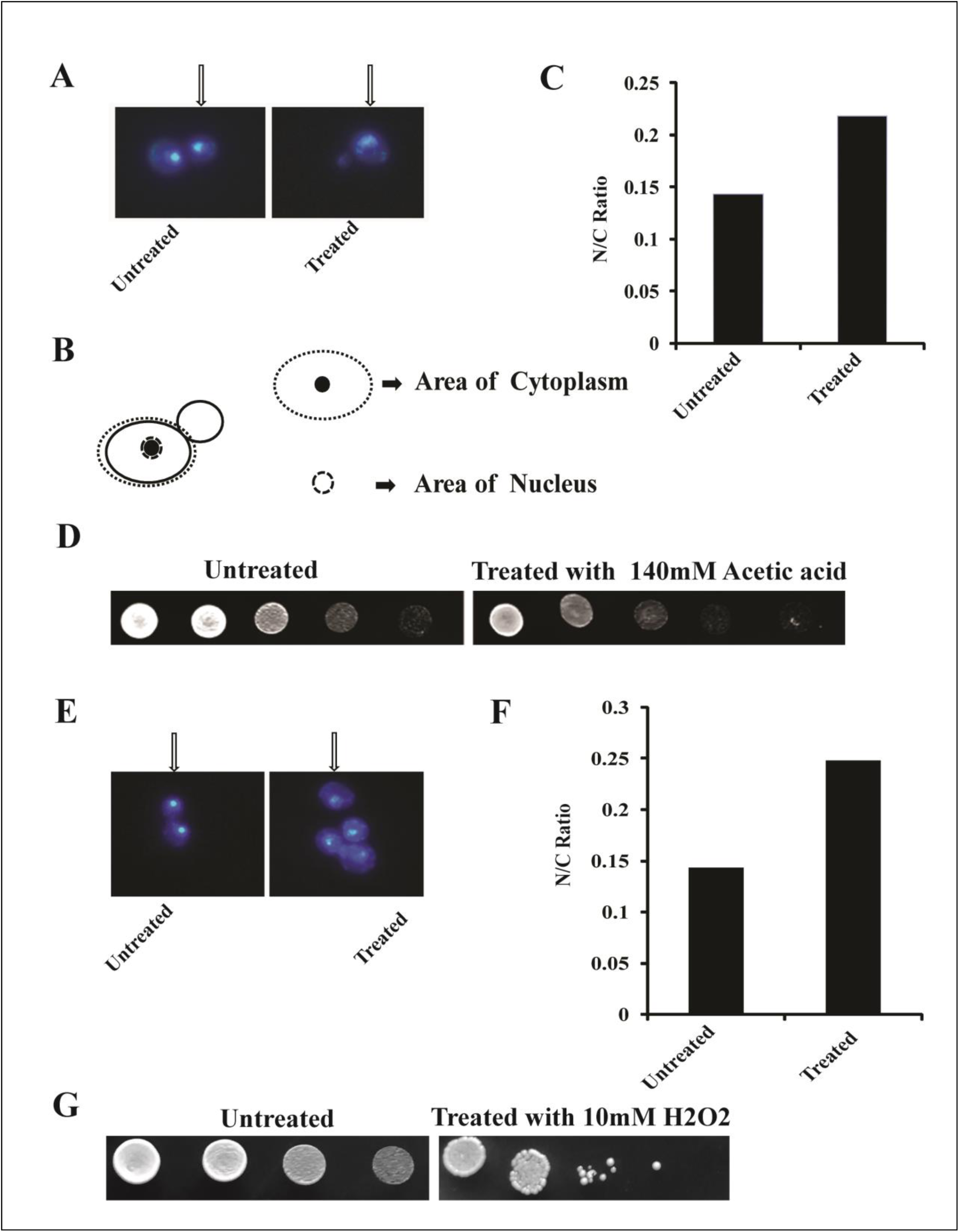
The qualitative and quantitative features of apoptosis as increase in nucleus to cytoplasm ratio in *S. cerevisiae.* **A, E.** Representative images of DAPI stained, *S. cerevisiae*, BY4741 cells before and after treatment of the apoptosis inducing concentration of acetic acid (140mM) and hydrogen peroxide (10mM) for 200 minutes respectively, arrow indicating the nuclear fragmentation with increased nuclear area in comparison to untreated cells, where nucleus is compact. **B.** Schematics indicating the scheme of quantification of area of nucleus and cytoplasm in the yeast cells. The average was calculated from the area of nucleus and cytoplasm of 25 cells from both untreated and treated cells and ratio of nucleus to cytoplasm was drawn. **C, F.** Bar diagrams showing the nucleus to cytoplasm ratio of the untreated and treated cells with acetic acid and hydrogen peroxide indicating a two fold increase in the nuclear to cytoplasm ratio in the apoptotic cells. **D, G.** Spot assay for comparative growth analysis of untreated and treated cells, indicating growth inhibition due to apoptosis of the treated cells. The cells overcome the oxidative stress mediated by hydrogen peroxide more rapidly than acetic acid as indicated by better growth in case of hydrogen peroxide treated cells.

**Figure 2.**
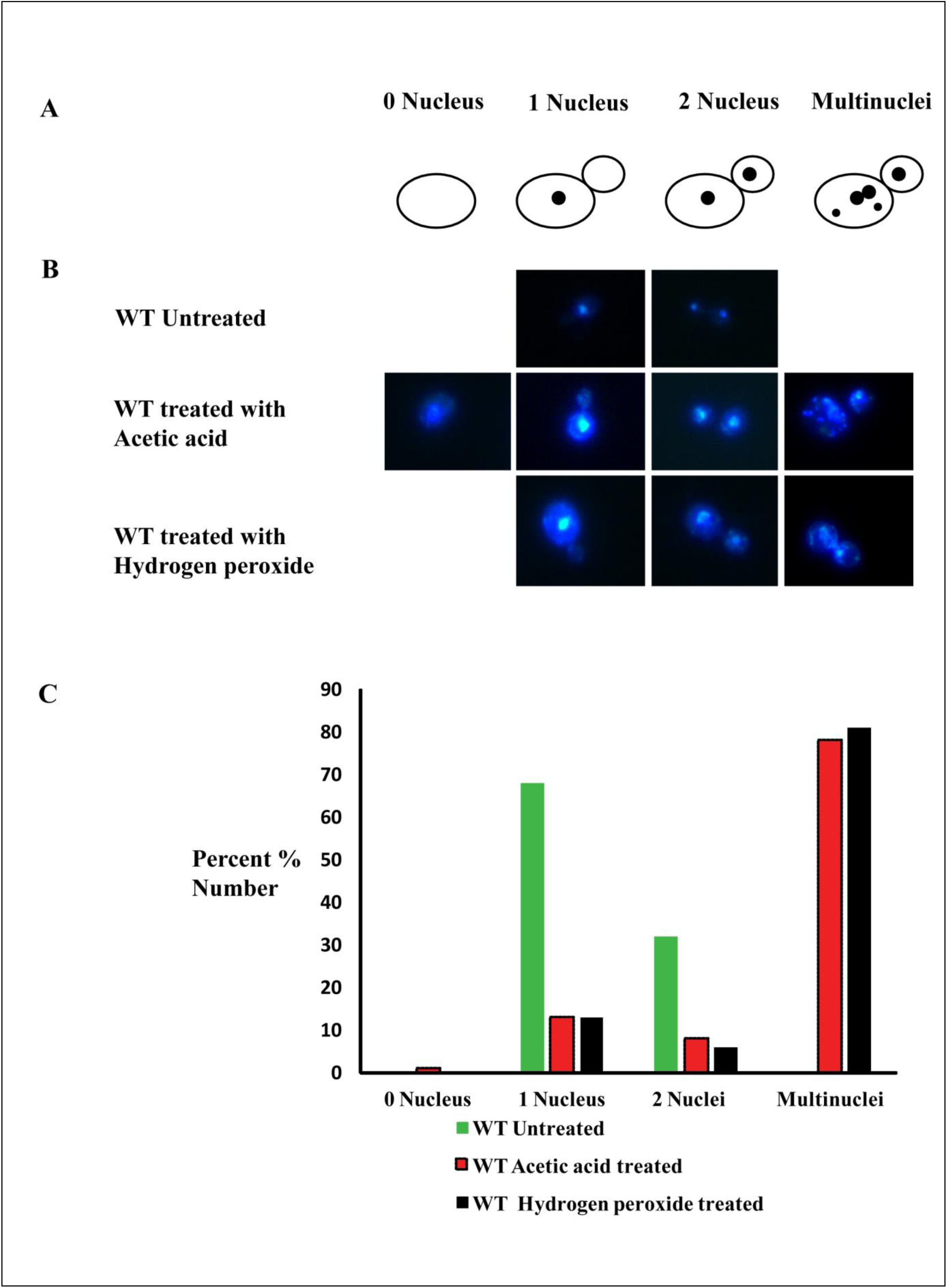
Increased multi nuclei as apoptotic feature of cells treated with apoptosis inducing concentration of acetic acid and hydrogen peroxide. **A.** Schematics of the nucleus status in cells, cells with no nucleus, 1 nucleus, 2 nuclei and multi nuclei **B.** Images of DAPI stained WT cells untreated and treated with acetic acid (140mM) and hydrogen peroxide (10mM) for 200 minutes indicating the 1 nucleus, two nuclei and multi nuclei phenotype. **C.** Bar diagram indicating the percent cells with 1 nucleus, 2 nuclei, and multi nuclei phenotype in untreated and treated WT cells.

The quantification of multi nuclei phenotype due to nuclear migration defects was first reported by (Brachat *et al.* 1998) while studying the spindle pole body defects in the *cnm67Δ1* and wild type CEN.PK2 strains. The faithful distribution of the duplicated nuclei is important for even distribution of genetic material from mother to daughter cell in eukaryotes. After duplication of genetic material the migration of nuclei in *S. cerevisiae* requires two major steps first, nucleus moves closer to the bud neck region second, insertion of the separating nucleus to the daughter cells during anaphase (Yeh *et al.* 1995; DeZwaan *et al.* 1997). The process of nuclear migration is dependent on the cytoplasmic microtubules and mutants of tubulin affect the nuclear migration leading to abnormal nuclear division in mother cells resulting accumulation of two or more nuclei (Huffaker *et al.* 1988; Palmer *et al.* 1992; Sullivan and Huffaker 1992). Here in this study we investigated the impact of apoptosis inducing doses of acetic acid and hydrogen peroxide on the nuclear migration in the WT cell (BY4741) after DAPI staining and characterized the nuclear migration phenotype as 0 nuclei, 1 nucleus, 2 nuclei and more than 3 nuclei (**Figure 2A**) as reported earlier (Brachat *et al.* 1998). The one and two nuclei are the normal state of the cells without any defects however cell showing the more than 3 nuclei can be termed as apoptotic cells (**Figure 2B**). The nuclear DNA fragmentation is the last stages of the apoptosis and it can be observed by DAPI staining in the *S. cerevisiae* cells if treated with the apoptosis inducing doses of the acetic acid. In this assay system when we compared the WT cells treated with the acetic acid and hydrogen peroxide there was dramatic increase of cells with the multi nuclei phenotype and nearly 70 % and above cells showed the multi nuclei phenotype (**Figure 2C**) in comparison to untreated cells. It is interesting to note that the cells with the multi nuclei may be the results of nuclear DNA fragmentation due to the stress caused by acetic acid and hydrogen peroxide. Based on the results, after the treatment of the acetic acid and hydrogen peroxide and combined observations on multi nuclei phenotype and nucleus to cytoplasm ratio indicate that these two parameters can be studied as apoptosis marker in *S. cerevisiae.* In the end we propose that the apoptosis can be studied both qualitatively and quantitatively as mentioned in the present study in *S. cerevisiae* and similar organisms.

## Acknowledgment

The authors would like to thank to Dr. Deepak Sharma, IMTECH, Dr. Ravi Manjithya, JNCASR, Dr. Jitendra Thakur, NIPGR, New Delhi, India for strains and Dr. Fayaz Malik, IIIM, Jammu & Kashmir for DAPI Stain.

## Compliance with Ethical Standards

Authors declares no conflict of interest.

## Ethical Approval

This article does not contain any studies with human participants performed by any of the authors.

## Author’s contributions

NKB conceived and directed the study and wrote the paper with all the authors. HS, MP, MS conducted the experiments. All the authors analysed the data, reviewed the results, and approved the final version of manuscript.

